# Regulation of Kidney Mitochondrial Function by Caloric Restriction

**DOI:** 10.1101/2021.12.29.474441

**Authors:** Julian D. C. Serna, Andressa G. Amaral, Camille C. Caldeira da Silva, Ana C. Munhoz, Sergio L. Menezes-Filho, Alicia J. Kowaltowski

## Abstract

Caloric restriction (CR) prevents obesity, promotes healthy aging, and increases resilience against several pathological stimuli in laboratory rodents. At the mitochondrial level, protection promoted by CR in the brain and liver is related to higher calcium uptake rates and capacities, avoiding Ca^2+^-induced mitochondrial permeability transition. Dietary restriction has also been shown to increase kidney resistance against damaging stimuli such as ischemia/reperfusion, but if these effects are related to similar mitochondrial adaptations had not yet been uncovered. Here, we characterized changes in mitochondrial function in response to six months of CR in rats, measuring bioenergetic parameters, redox balance and calcium homeostasis. CR promoted an increase in mitochondrial oxygen consumption rates under non-phosphorylating and uncoupled conditions. While CR prevents mitochondrial reactive oxygen species production in many tissues, in kidney we found that mitochondrial H_2_O_2_ release was enhanced, although levels of carbonylated proteins and methionine sulfoxide were unchanged. Surprisingly, and opposite to the effects observed in brain and liver, mitochondria from CR animals are more prone to Ca^2+^-induced mitochondrial permeability transition. CR mitochondria also displayed higher calcium uptake rates, which were not accompanied by changes in calcium efflux rates, nor related to altered inner mitochondrial membrane potentials or the amounts of the mitochondrial calcium uniporter (MCU). Instead, increased mitochondrial calcium uptake rates in CR kidneys correlate with a loss of MICU2, an MCU modulator. Interestingly, MICU2 is also modulated by CR in liver, suggesting it has a broader diet-sensitive regulatory role controlling mitochondrial calcium homeostasis. Together, our results highlight the organ-specific bioenergetic, redox, and ionic transport effects of CR. Specifically, we describe the regulation of the expression of MICU2 and its effects on mitochondrial calcium transport as a novel and interesting aspect of the metabolic responses to dietary interventions.

## Introduction

Kidneys are essential for homeostatic regulation of the volume and composition of body fluids. These activities rely on the active transport of ions and molecules in nephrons (1,2), an energetically demanding activity maintained mainly through aerobic metabolism, as reflected by their high mitochondrial content and oxygen consumption rates (3,4). Oxidative phosphorylation (OxPhos) is finely coordinated with immediate cell energetic needs, not only by canonical modulation of ATP synthase by ADP availability, but also through activation of electron transport and mitochondrial matrix enzymes by Ca^2+^ ions (5).

Indeed, mitochondria take up Ca^2+^ in a highly regulated fashion, allowing these organelles to integrate their activity with spatiotemporal alterations in cytoplasmic Ca^2+^ concentrations (6). At the cellular level, these Ca^2+^ signals control processes as diverse as muscle contraction, proliferation, and hormone secretion, all of which require increased energy input. In the mitochondrial matrix, Ca^2+^ directly or indirectly activates several dehydrogenases of the tricarboxylic acid cycle. At the inner mitochondrial membrane, Ca^2+^ activates proteins involved in substrate transport (7). Overall, this results in increased mitochondrial ATP synthesis capacity (5). However, these desirable effects of mitochondrial Ca^2+^ uptake can also have a downside: excessive Ca^2+^ may favor the production reactive oxygen species such as superoxide radicals and H_2_O_2_, and induce oxidative loss of inner mitochondrial membrane integrity, known as the mitochondrial permeability transition (mPT) (8).

Mitochondrial Ca^2+^ uptake occurs through the MCU complex (9), which in metazoans is formed, mainly, by the *mitochondrial calcium uniporter* (MCU), *essential MCU regulator* (EMRE), and *mitochondrial calcium uptake proteins* 1 and 2 (MICU1, and MICU2). An MCU tetramer forms an electronegative path for Ca^2+^ diffusion through the inner mitochondrial membrane (10,11). EMRE, in a proportion 1:1 with MCU, maintains the channel in an active conductive state (12,13). MICU1 and MICU2 (as well as MICU3, in muscle and the central nervous system) regulate the kinetics and threshold of Ca^2+^ uptake in a non-redundant fashion (14–16). MICU2 forms an obligate heterodimer with MICU1 (which itself can form homodimers under certain conditions). MICU1 maintains the MCU complex closed at low Ca^2+^ levels, and regulates its cooperative calcium transport behavior. The specific role of MICU2 is less understood, but several studies suggest it negatively regulates Ca^2+^ uptake (16,17). The structure of the MCU complex is not fixed, and several variations of it have been described. Indeed, alterations of the proportion of MICU1 or 2 to MCU were shown to control tissue-specific decoding of cytoplasmic Ca^2+^ signals by mitochondria (18). MICU composition has shown to be regulated in several pathological contexts such as diabetes, cancer, and polycystic disease (19–21).

Acute kidney injury is one of the main causes of renal failure, often occurring due to an ischemic event (22). Cytosolic Ca^2+^ overload has a central role in the pathophysiology of cell injury during ischemia/reperfusion, conditions in which excessive mitochondrial Ca^2+^ uptake may result in mitochondrial damage through induction of mPT, as discussed above. mPT involves an increase in the permeability of the inner mitochondrial membrane, disrupting ion balance and hampering oxidative phosphorylation; it may eventually culminate in cell death (8,23,24). Susceptibility to mPT is modulated by several factors. For example, aging and conditions that promote unhealthy aging typically make mitochondria more prone to mPT (25–27). In contrast, caloric restriction (CR), a dietary intervention that encompasses a decrease in energy intake (without malnutrition) and is well known to increase lifespans and improve healthspans, protects against Ca^2+^-induced mPT in the liver and the brain (28–30).

The effects of CR on mitochondrial Ca^2+^ transport are, however, organ-specific (31), and have not been explored in kidneys to date. Kidneys are known to be protected from damage induced by ischemia/reperfusion in animals submitted to dietary restriction or fasting (32,33). Furthermore, these protective effects can be mimicked by cyclosporin A (CsA) a well-known inhibitor of mPT, suggesting that dietarily-restricted kidneys may display protection against mitochondrial Ca^2+^-induced damage.

Here, we investigated the effects of CR on mitochondrial function, and surprisingly found that CR regulates Ca^2+^ uptake in kidney mitochondria, but in the opposite manner than that observed in brain and liver, decreasing maximum uptake capacity and favoring mPT. Our results reinforce prior data indicating that dietary interventions affect mitochondrial Ca^2+^ uptake in an organ-specific manner, and correlate the regulation of Ca^2+^ uptake in mitochondria by diets to changes in the composition of the MCU complex.

## Materials and Methods

### Animal care and caloric restriction

Male Sprague Dawley rats (NTac: SD) were bred and lodged at the local Animal Facility (*Instituto de Química, Universidade de São Paulo*). All experiments were approved by the ethics committee for animal care and use and performed following NIH guidelines. Rats were housed in plastic ventilated cages (49 cm x 34 cm x 16 cm) under specific pathogen free conditions at constant temperature (22 ± 2°C) and relative air humidity (50 ± 10%) The photoperiod was 12 h light/12 h dark, with the light period beginning at 6:00. Soon after they reached adulthood (~12 weeks old) the dietary intervention was initiated, and maintained for a total of 6 months. Animals were randomly separated into an *ad-libitum* (AL) group, with free access to a standard AIM93G diet, or a caloric restriction (CR) group, with a reduction in 40% of the amount of food consumed by AL rats. To avoid malnutrition, CR rats were fed with an AIM93G diet supplemented with vitamins and minerals to the same levels as AL animals consumed (34). Details regarding the CR protocol and its effects can be consulted in our previous publications (29–31). Prior to the experiments, animals were fasted for 10-12 h, beginning at 19:00. Animals were then deeply anesthetized (1 mL · Kg^−1^ ketamine, 0.6 mL · Kg^−1^ xylazine, and 0.5 mL · Kg^−1^ acepromazine, subcutaneously) and euthanized by cardiac puncture, followed by a diaphragm incision.

### Mitochondrial isolation

Kidney mitochondria were isolated through differential centrifugation as described by Tahara et al. (35), with some modifications. All steps were carried out at 4°C, over ice. Kidneys were cut into small pieces (2-3 mm) and washed several times with 10 mM EDTA-PBS. The chopped organs were suspended in isolation buffer (0.3 M sucrose, 10 mM HEPES, 2 mM EGTA, 1 mM EDTA; pH = 7.2) supplemented with 0.2% BSA, and homogenized in an Elvehjem Potter with at least 5 strokes. The mixture was centrifuged at 800 g for 5 minutes, and the pellet was discharged. This step was performed twice to guarantee that mitochondrial preparations were not contaminated with blood. The supernatant was further centrifuged at 12,000 g for 10 min. The resulting pellet was washed, resuspended in isolation buffer and centrifuged, again, at 12,000 g for 10 min. The small fluffy layer present in all the pellets was included in the final preparation so as not to select mitochondrial populations in the different experimental groups. The protocols for the isolation of heart and liver mitochondria are detailed in Serna, et al. (31) and Menezes Filho, et al. (30). Protein concentration was determined using the Bradford method.

### Ca^2+^ uptake assays

Kidney mitochondria-enriched fractions (500 μg) were incubated in 2 mL of experimental buffer (125 mM sucrose, 65 mM KCl, 10 mM Hepes, 2 mM K^+^ phosphate, 2 mM MgCl2, and 0.2% BSA; pH was adjusted to 7.2 with KOH), in the presence of 0.1 μM Calcium Green 5N, 1 mM succinate and 2 μM rotenone. Calcium Green 5N fluorescence (excitation and emission wavelengths of 506 nm and 532 nm, respectively) was measured with a F4500 Hitachi Fluorimeter at 37°C, under constant stirring (29–31). The equation [Ca^2+^] = K_d_·(F-F_min_)/(F_max_-F) was employed to determine the relationship between fluorescence (F) and [Ca^2+^] concentrations. The K_d_ value was determined as the value at which the change in fluorescence (ΔF) after each calcium addition can be matched by a 10 μM change in calcium concentration (Δ[Ca^2+^]). Maximal (F_max_) and minimal (F_min_) fluorescence was determined at the end of each trace using 100 μL of 100 mM Ca^2+^ and 100 mM EGTA solutions, respectively. To determine calcium retention capacities (in nmol Ca^2+^· mg protein^−1^), several additions of 10 μM Ca^2+^ were made until mPT occurred. These traces were performed with and without 10 μM Cyclosporin A (CsA), as indicated. Calcium uptake rates (in nmol Ca^2+^. mg protein^−1^. s^−1^) were determined as the slope of the linear portion at the beginning of the first calcium addition.

### Ca^2+^ efflux assays

Briefly, 500μg of mitochondrial protein were loaded with 60 nmol Ca^2+^, as described for the uptake assays. Soon after complete uptake, 10 μM Ruthenium Red (RuR), an MCU inhibitor, was added. Ca^2+^ efflux was then stimulated with 20 mM Na^+^ to measure mitochondrial sodium calcium exchanger (NCLX)-dependent Ca^2+^-efflux (36). Calcium efflux rates were measured as the slope of the linear portion at the beginning of the traces after the addition of RuR (basal) or Na^+^ (stimulated).

### Mitochondrial oxygen consumption

Mitochondrial respiration was assessed using a high-resolution oxygraph (Oroboros). Mitochondria (100 μg) was suspended in 2 mL of respiration buffer (experimental buffer supplemented with 1 mM EGTA). Mitochondrial activity was modulated through the sequential addition of 1 mM succinate and 2 μM rotenone (state 2), 1 mM ADP (state 3), 1 μM oligomycin (state 4) and 1.25 μM CCCP (state 3U). Respiratory control ratios (RCR) were determined as the ratio between oxygen consumption rates under state 3 and state 4.

### Mitochondrial membrane potential measurements

Mitochondrial membrane potentials (ΔΨ) were determined by measuring changes in the fluorescence of safranin O, in quenching mode (37). Kidney mitochondria (75 μg) were incubated in 2 mL of respiration buffer in the presence of 5 μM Safranin O, 1 mM succinate, and 1 μM rotenone. Fluorescence was followed in an F4500 Hitachi Fluorimeter at excitation and emission wavelengths of 485 nm and 586 nm, respectively, under constant stirring, at 37°C. A calibration curve was employed to establish a relationship between ΔΨ and fluorescence (38). Briefly, for calibration, ΔΨ was clamped at different values using valinomycin (ionophore) and different known KCl concentrations. At each point, safranin O fluorescence was determined and the calibration curve was constructed.

### Mitochondrial hydrogen peroxide release

Mitochondrial H_2_O_2_ release was followed through horseradish peroxidase (5U/mL)-catalyzed oxidation of 25 μM Amplex Red (39). 75 μg of kidney mitochondria were added to 2 mL of experimental buffer in the presence of 1 mM succinate and 1 μM rotenone, under constant stirring, at 37°C. The oxidation of Amplex Red generates a fluorescent compound known as resorufin, followed with an F2500 Hitachi Fluorimeter at excitation and emission wavelength of 563 nm and 587 nm, respectively. To establish a relationship between resorufin fluorescence and H_2_O_2_ concentrations, a calibration curve was constructed under the same experimental conditions in the absence of mitochondria, by sequentially adding known quantities of H_2_O_2_.

### Complex I (NADH: ubiquinone oxidoreductase) specific activity

The specific enzymatic activity of mitochondrial Complex I was measured spectrophotometrically in isolated mitochondria through the decrease in NADH (40); the original protocol was adapted for 96 well plates. Briefly, mitochondria were incubated in hypotonic Tris buffer (10 mM Tris, pH = 7.6) and subjected to three freeze-thaw cycles. This process guarantees that the inner mitochondrial membrane is completely disrupted and substrates are freely available. The disrupted mitochondrial suspension (1 μg) was incubated in reaction buffer for complex I activity (50 mM K^+^ phosphate buffer, 3 mg · mL^−1^ fatty acid-free BSA, 300 μM KCN and 100 μM NADH; pH = 7.5). The reaction was initiated with 60 μM ubiquinone1. The absorbance of the sample at 340 nm (ε = 6.2 mM^−1^ · cm^−1^) decreases with the activity of Complex I. NADH disappearance catalyzed by enzymes other than Complex I was estimated using 10 μM rotenone, a Complex I inhibitor. The reaction was followed for at least 5 minutes at 37°C. Enzyme kinetics were analyzed under initial velocity conditions. Data were normalized to the total mitochondrial protein and to specific Complex I activity of AL controls.

### Complexes I+III (NADH cytochrome c oxidoreductase)

Specific enzymatic activity of Complexes I + III was measured through the reduction of cytochrome c (40), which absorbs light more strongly at λ = 550 nm (ε = 19.5 mM^−1^ · cm^−1^) than the oxidized form. Mitochondria were incubated in hypotonic Tris buffer, as described above, and subjected to three freeze-thaw cycles. 2 μg of the mitochondrial suspension were incubated in reaction buffer (50 mM K^+^ phosphate buffer, 1 mg · mL^−1^ fatty acid-free BSA, 300 μM KCN and 50 μM cytochrome c; pH = 7.5). The reaction was initiated with 200 μM NADH. Enzymatic kinetics were analyzed under initial velocity conditions. Data were normalized to the specific Complex I + III activity of AL controls.

### Complex II (succinate: ubiquinone oxidoreductase) specific activity

Complex II enzymatic activity was measured spectrophotometrically in isolated mitochondria through the reduction of DCPIP (2,6-dichlorophenolindophenol), which loses its color when reduced (40). Mitochondria were incubated in hypotonic Tris buffer (see above) and subjected to three freeze-thaw cycles. Then, 0.5 μg of the mitochondrial suspension was incubated in Complex II reaction buffer (25 mM K^+^ phosphate buffer, 1 mg · mL^−1^ fatty acid-free BSA, 300 μM KCN, 20 mM succinate, and 80 μM DCPIP; pH = 7.5) for 10 minutes at 37°C. The reaction was initiated with 50 μM decylubiquinone (DUB) and followed for 10 min. Normally, Complex II oxidizes succinate and the electrons are transported to Complex III through the electron carrier ubiquinone. DUB was employed as an electron carrier and DCPIP as the final electron acceptor. The rate of DCPIP reduction can be followed by a decrease in the absorbance of the sample at 600 nm (ε = 19.1 mM^−1^ · cm^−1^). Enzymatic kinetics were analyzed under initial velocity conditions. Data was normalized to the specific Complex II activity of AL controls.

### Complexes II + III (succinate cytochrome c oxidoreductase)

Complex II + III-specific activities was measured through the reduction of cytochrome c (40) followed at λ = 550 nm (ε = 19.5 mM^−1^ · cm^−1^). Mitochondria were incubated in hypotonic Tris buffer and subjected to three freeze-thaw cycles. 2 μg of the mitochondrial suspension were incubated in reaction buffer (100 mM K^+^ phosphate buffer, 300 μM KCN, and 10 mM succinate; pH 7.5) for 10 min. The reaction was initiated 50 μM cytochrome c. The reaction was followed for at least 5 minutes at 37°C. Enzymatic kinetics were analyzed under initial velocity conditions. Data were normalized to the specific activity of AL controls.

### Western blots

After isolation, a fraction of the mitochondrial preparations was collected and frozen at −80°C. Kidneys were homogenized in T-PER™ (Tissue Protein Extraction Reagent, Thermo Scientific™ 78510) with Protease and Phosphatase Inhibitor Cocktail (Halt Protease & Phosphatase Single-Use Inhibitor Cocktail, Thermo Scientific™ 78442) using a polytron, centrifuged at 10,000 g for 10 min at 4°C, and the supernatant was collected and stored at −80°C. Protein concentration was measured through the Bicinchoninic Acid protein assay (Pierce™ BCA Protein Assay Kit, Thermo Scientific™ 23225). Equal amounts of proteins extracts (20-30 μg for mitochondrial samples or 40 μg for total kidney homogenates) were loaded in acrylamide gels. Sample preparation and electrophoresis (41) conditions were detailed in Table 1. Proteins were electrotransferred (42) in a wet system at 100 V for 1-2 h, onto a nitrocellulose or PVDF membrane. After transfer, membranes were stained with Ponceau Red and washed with Milli-Q water. Ponceau stains were used as protein loading controls in order to avoid bias promoted by specific protein controls. Membranes were blocked in 5% defatted milk diluted in TBST (20 mM Tris, 150 mM NaCl, 0.1% Tween 20, pH 7.4) for 1 h at room temperature. All primary antibodies (Table 2) were diluted in a BSA solution (5% in TBST) and were incubated overnight at 4°C. After washing in TBST, each membrane was incubated during 1 h with an appropriate secondary antibody (Table 2). Images were processed and quantified in Image J. Target protein signal was normalized by total lane signal of the membrane after Ponceau staining. Data were presented as fold change relative to AL.

**Table 1.**
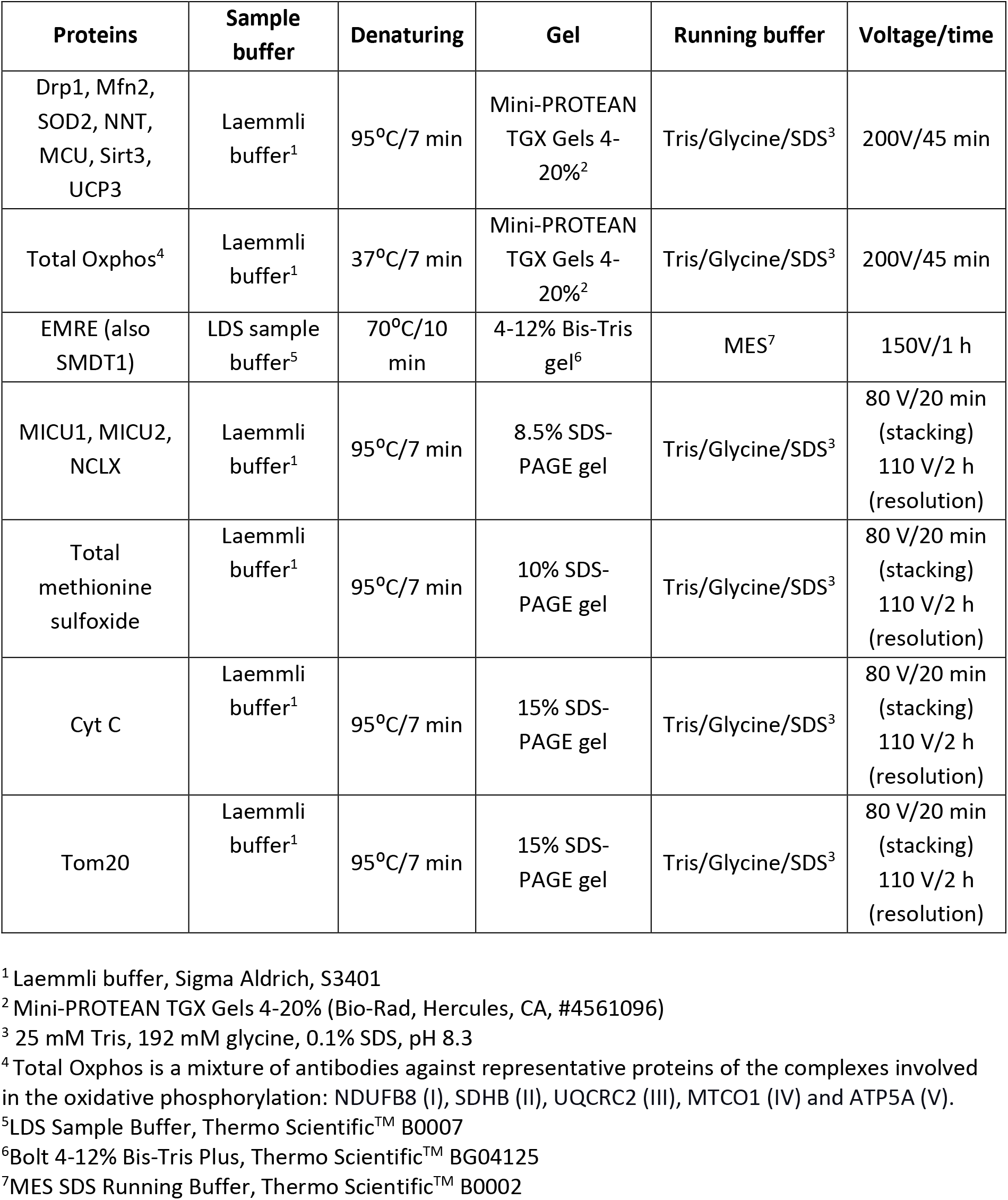
Sample preparation and run conditions.

**Table 2.**
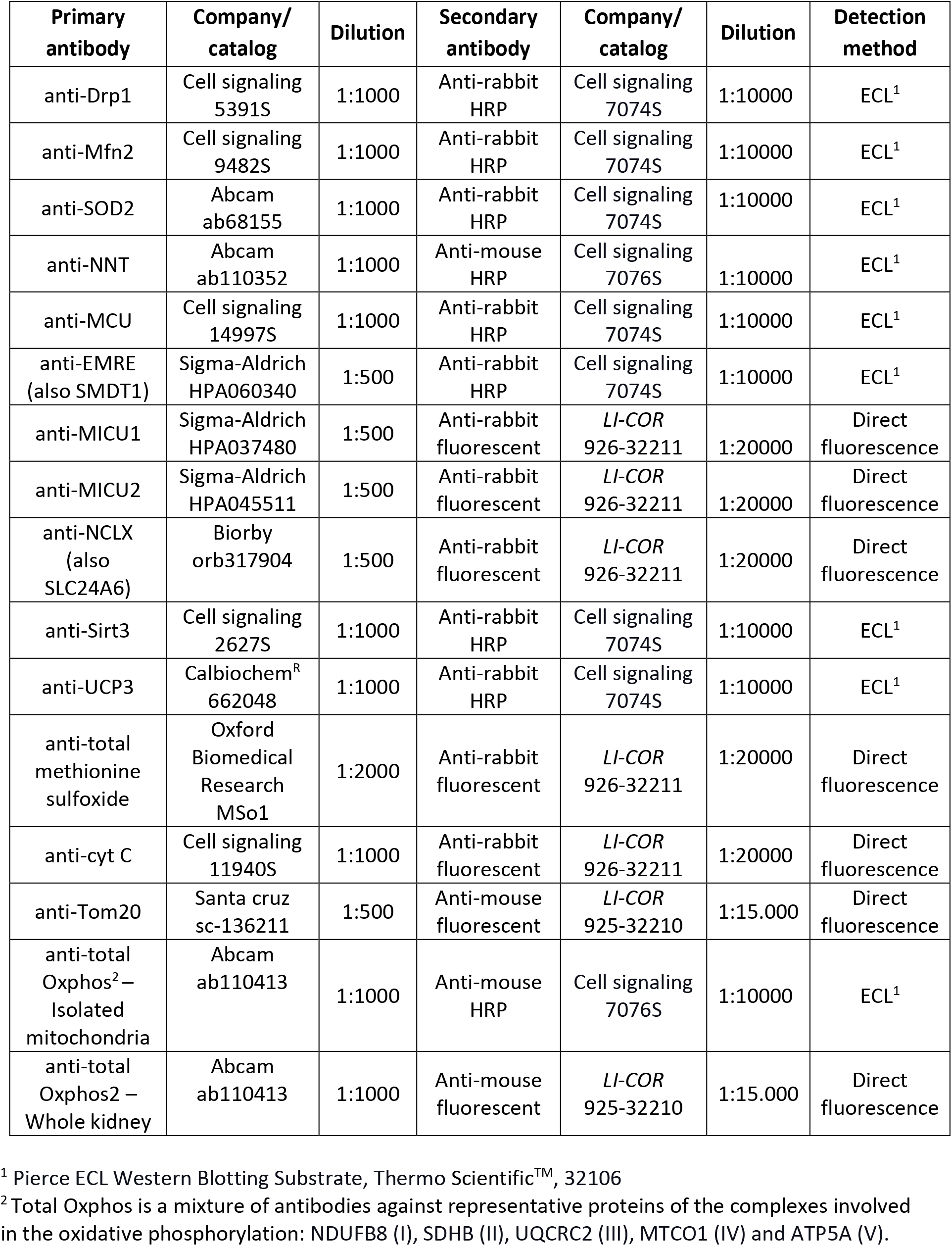
Primary and secondary antibodies.

| Primary antibody | Company/<br>catalog | Dilution | Secondary antibody | Company/<br>catalog | Dilution | Detection method |
| --- | --- | --- | --- | --- | --- | --- |
| anti-Drp1 | Cell signaling<br>5391S | 1:1000 | Anti-rabbit<br>HRP | Cell signaling<br>7074S | 1:10000 | ECL<sup>1</sup> |
| anti-Mfn2 | Cell signaling<br>9482S | 1:1000 | Anti-rabbit<br>HRP | Cell signaling<br>7074S | 1:10000 | ECL<sup>1</sup> |
| anti-SOD2 | Abcam<br>ab68155 | 1:1000 | Anti-rabbit<br>HRP | Cell signaling<br>7074S | 1:10000 | ECL<sup>1</sup> |
| anti-NNT | Abcam<br>ab110352 | 1:1000 | Anti-mouse<br>HRP | Cell signaling<br>7076S | 1:10000 | ECL<sup>1</sup> |
| anti-MCU | Cell signaling<br>14997S | 1:1000 | Anti-rabbit<br>HRP | Cell signaling<br>7074S | 1:10000 | ECL<sup>1</sup> |
| anti-EMRE<br>(also SMDT1) | Sigma-Aldrich<br>HPA060340 | 1:500 | Anti-rabbit<br>HRP | Cell signaling<br>7074S | 1:10000 | ECL<sup>1</sup> |
| anti-MICU1 | Sigma-Aldrich<br>HPA037480 | 1:500 | Anti-rabbit<br>fluorescent | *LI-COR*<br>926-32211 | 1:20000 | Direct<br>fluorescence |
| anti-MICU2 | Sigma-Aldrich<br>HPA045511 | 1:500 | Anti-rabbit<br>fluorescent | *LI-COR*<br>926-32211 | 1:20000 | Direct<br>fluorescence |
| anti-NCLX<br>(also<br>SLC24A6) | Biorby<br>orb317904 | 1:500 | Anti-rabbit<br>fluorescent | *LI-COR*<br>926-32211 | 1:20000 | Direct<br>fluorescence |
| anti-Sirt3 | Cell signaling<br>2627S | 1:1000 | Anti-rabbit<br>HRP | Cell signaling<br>7074S | 1:10000 | ECL<sup>1</sup> |
| anti-UCP3 | Calbiochem<sup>R</sup><br>662048 | 1:1000 | Anti-rabbit<br>HRP | Cell signaling<br>7074S | 1:10000 | ECL<sup>1</sup> |
| anti-total<br>methionine<br>sulfoxide | Oxford<br>Biomedical<br>Research<br>MSo1 | 1:2000 | Anti-rabbit<br>fluorescent | *LI-COR*<br>926-32211 | 1:20000 | Direct<br>fluorescence |
| anti-cyt C | Cell signaling<br>11940S | 1:1000 | Anti-rabbit<br>fluorescent | *LI-COR*<br>926-32211 | 1:20000 | Direct<br>fluorescence |
| anti-Tom20 | Santa cruz<br>sc-136211 | 1:500 | Anti-mouse<br>fluorescent | *LI-COR*<br>925-32210 | 1:15.000 | Direct<br>fluorescence |
| anti-total<br>Oxphos<sup>2</sup> –<br>Isolated<br>mitochondria | Abcam<br>ab110413 | 1:1000 | Anti-mouse<br>HRP | Cell signaling<br>7076S | 1:10000 | ECL<sup>1</sup> |
| anti-total<br>Oxphos2 –<br>Whole kidney | Abcam<br>ab110413 | 1:1000 | Anti-mouse<br>fluorescent | *LI-COR*<br>925-32210 | 1:15.000 | Direct<br>fluorescence |

<sup>1</sup> Pierce ECL Western Blotting Substrate, Thermo Scientific™, 32106
<sup>2</sup> Total Oxphos is a mixture of antibodies against representative proteins of the complexes involved in the oxidative phosphorylation: NDUFB8 (I), SDHB (II), UQCRC2 (III), MTCO1 (IV) and ATP5A (V).

## Statistical analysis

Statistical analysis was performed using GraphPad Prism Software 4.0. Most data were analyzed using a two tailed unpaired t-tests. A matched one-way ANOVA with Holm-Šídák post hoc test was employed in Fig. 5 for multiple comparisons. A p value ≤ 0.05 was considered statistically significant.

## Results

While the effects of caloric restriction (CR) on energy metabolism in many organs have been explored in a detailed manner, little attention has been focused on the metabolic effects of this dietary intervention in the kidney. In order to assess the renal metabolic effects of CR, rats were submitted for 6 months to either an *ad libitum* (AL) diet or a CR protocol in which food intake was decreased daily by 40%, without changes in micronutrient intake. Restricted dietary protocols are often found to change mitochondrial mass in specific tissues (43–47), so we evaluated possible changes in mitochondrial mass in whole kidney homogenates. Figure 1 shows that the quantities of five representative proteins that participate in oxidative phosphorylation (complexes I-V and cytochrome c) as well as Tom20, which participates in protein import, were unchanged in AL versus CR kidney homogenates. This indicates that mitochondrial mass is unaltered by CR.

**Figure 1.**
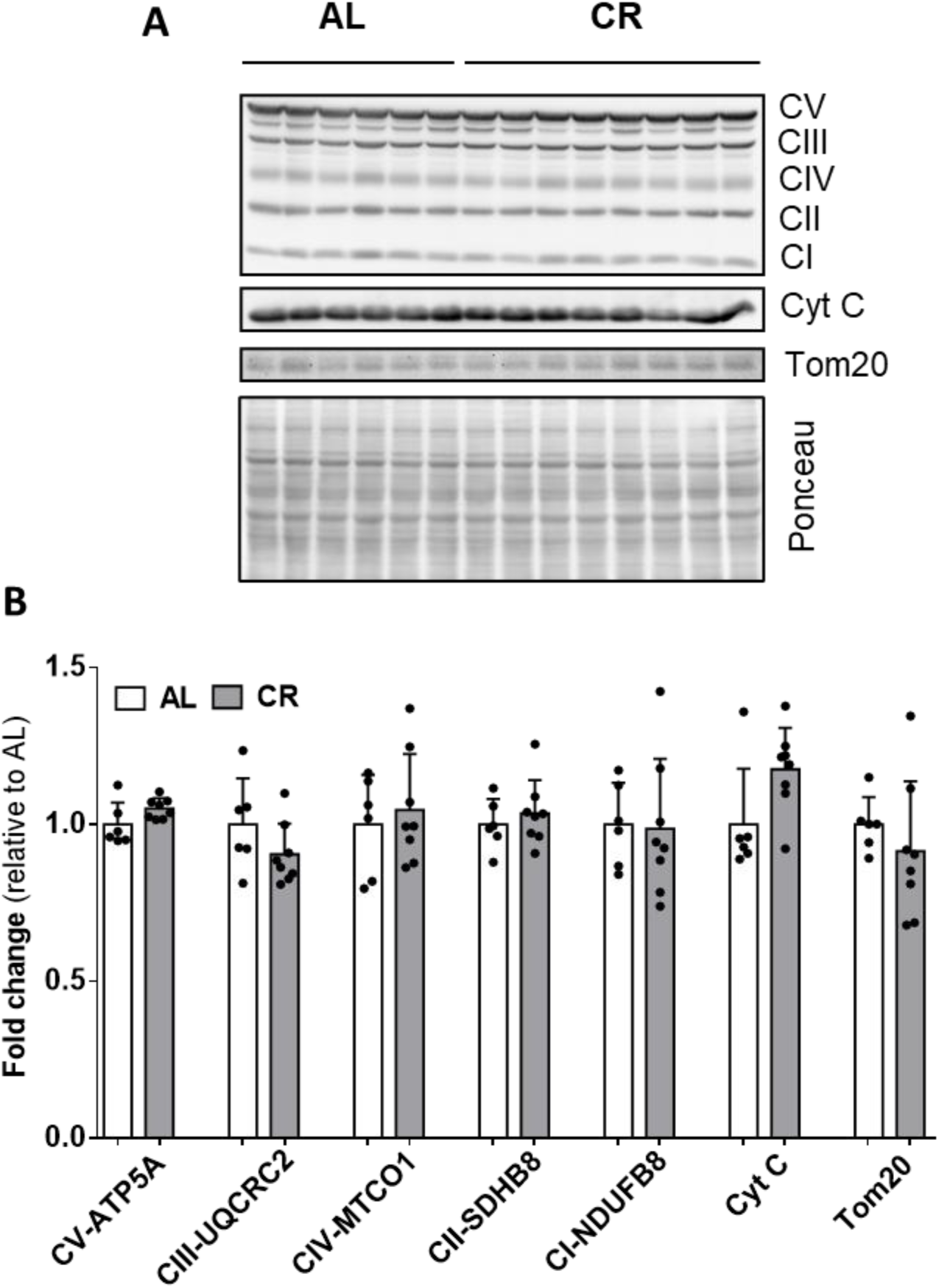
Kidney mitochondrial mass is not altered by caloric restriction. Changes in mitochondrial mass were probed through western blot analysis of mitochondrial proteins in whole kidney homogenates. **A.** Representative blot images for electron transport chain complex proteins (NDUFB8, SDHB, UQCRC2 and MTCO1), ATP synthase (ATP5A), Tom20 and cytochrome c (cyt c). **B.** Quantification relative to AL; the signal was normalized by the total Ponceau staining signal for each lane. Light and dark bars represent samples from AL and CR rats, respectively.

Although the quantities of mitochondrially-located proteins were similar in AL and CR kidneys, this does not necessarily reflect unaltered bioenergetic function. We thus characterized oxidative phosphorylation profiles of isolated mitochondria from AL and CR kidneys using high-resolution respirometry (Figure 2). Under conditions where only substrate and mitochondria were present (State 2), oxygen consumption rates were slightly, but significantly, higher in the CR group (Fig. 2A). When ATP synthesis was stimulated by adding ADP (State 3), no differences were observed among groups. In the presence of the ATP synthase inhibitor oligomycin A (state 4), and the uncoupler CCCP (state 3U), mitochondria from CR rats exhibited higher respiratory rates. Although changes in maximized electron transport (state 3U) and proton leak-mediated transport (state 4) were observed, respiratory control ratios (State 3/State4, Fig. 2B) were not significantly different.

**Figure 2.**
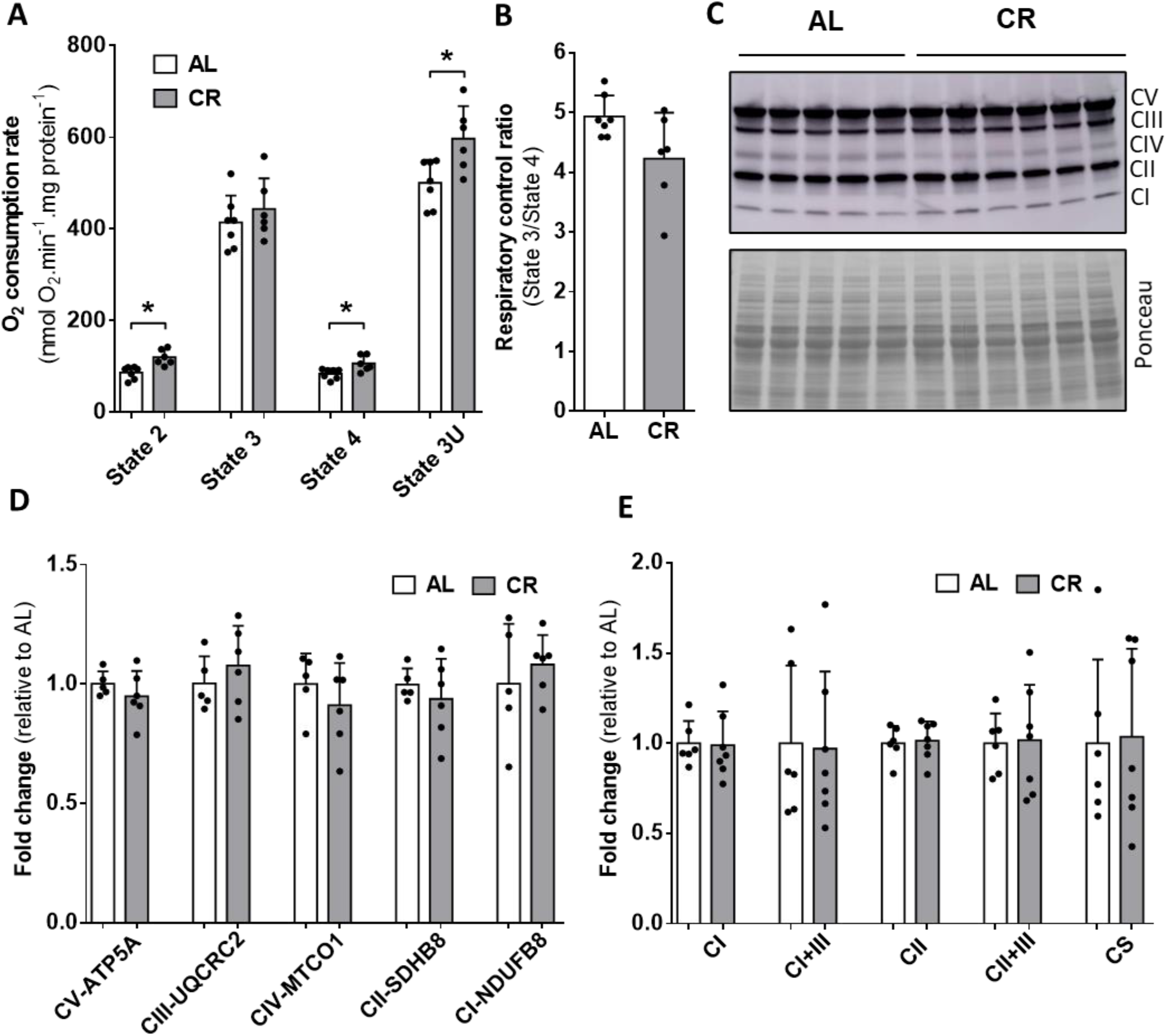
CR promotes subtle changes in kidney mitochondrial oxidative phosphorylation. **A.** Mitochondria were incubated as described in Methods, in the presence of 2 mM succinate plus 1 μM rotenone (state 2), 1 mM ADP (state 3), 1 μM oligomycin (state 4) and 1 μM CCCP (state 3U), added successively. Oxygen consumption rates were measured using high-resolution respirometry. **B.** Respiratory control ratios were calculated as the ratio between state 3 and state 4. **C.** The content of each complex of the electron transport chain (complexes I-IV) and the ATP synthase (complex V) was determined by the western blot analysis of the protein levels of NDUFB8 (I), SDHB (II), UQCRC2 (III), MTCO1 (IV) and ATP5A (V). **D.** Densitometric quantification for the blot signals, relative to AL. Protein was normalized in each lane by the total Ponceau stain. **E.** Respiratory complex enzymatic activities were determined spectrophotometrically for isolated complexes I and II, and combined complexes I+III and II+III. Light and dark bars represent AL and CR mitochondria, respectively. * p<0.05.

In order to attempt to uncover reasons for the increase in electron transport observed in Fig. 2A, we quantified the components of the electron transport chain and ATP synthase in isolated mitochondrial samples, an experimental condition in which nuanced differences not seen in whole tissue homogenates could be uncovered. However, we found no such changes (Figs. 2C and D). Higher respiratory rates in CR mitochondria could also result from higher respiratory complex activity, but no changes in enzymatic activity of complexes I and II was measured, and the combined activities of complexes I+III and II+III were also equal (Fig. 2E). Overall, these experiments show that CR promotes a small but significant increase in mitochondrial respiration under non-phosphorylating conditions, which is not associated with enhancement in the quantities or activities of respiratory complexes.

A potential reason for faster respiratory rates without alterations in mitochondrial respiratory chain component quantities or electron transport complex activities is a change in matrix and membrane enzymatic activities by Ca^2+^, which would require intact mitochondria to occur (5,6). Indeed, we have previously found that CR increases mitochondrial Ca^2+^ uptake in brain and liver mitochondria, although this effect is tissue-specific and not observed in cardiac or skeletal muscle (28–30). Prior data has also suggested that increased resistance to calcium-induced mPT may explain the higher resilience observed in kidneys from short-term dietarily-restricted (whole food restriction for 2-4 weeks) or fasted animals to damage-induced ischemia/reperfusion (32,33).

As the effects of CR on kidney mitochondria Ca^2+^ transport and mPT were not yet directly assessed in the literature, we investigated the possible effects of CR on isolated kidney mitochondrial Ca^2+^ transport properties (Figure 3). Mitochondria were suspended in an experimental media containing the extramitochondrial fluorescent probe Calcium Green-5N. The fluorescence of this molecule increases after calcium binding, which is proportional to extramitochondrial calcium concentrations. Fig. 3A shows a typical Ca^2+^ uptake trace: after each addition, at 200, 400, 600 and 800 s, a peak in the extramitochondrial Ca^2+^ concentration is observed. As the ion is taken up by mitochondria, fluorescence decreases. Each trace finished after the induction of the mitochondrial permeability transition (mPT), characterized by a loss of inner membrane impermeability and release of intramitochondrial Ca^2+^, thus increasing fluorescence. Calcium retention capacities were compared between AL and CR mitochondria (light and dark lines/bars, respectively) from traces such as Fig. 3A, and quantified in Fig. 3B. Surprisingly, and different from our previously published findings in brain and liver (29,30), CR significantly decreases Ca^2+^ uptake capacity in kidney mitochondria (Fig. 3B), an effect reversed by the mPT inhibitor CsA (Fig. 3C).

**Figure 3.**
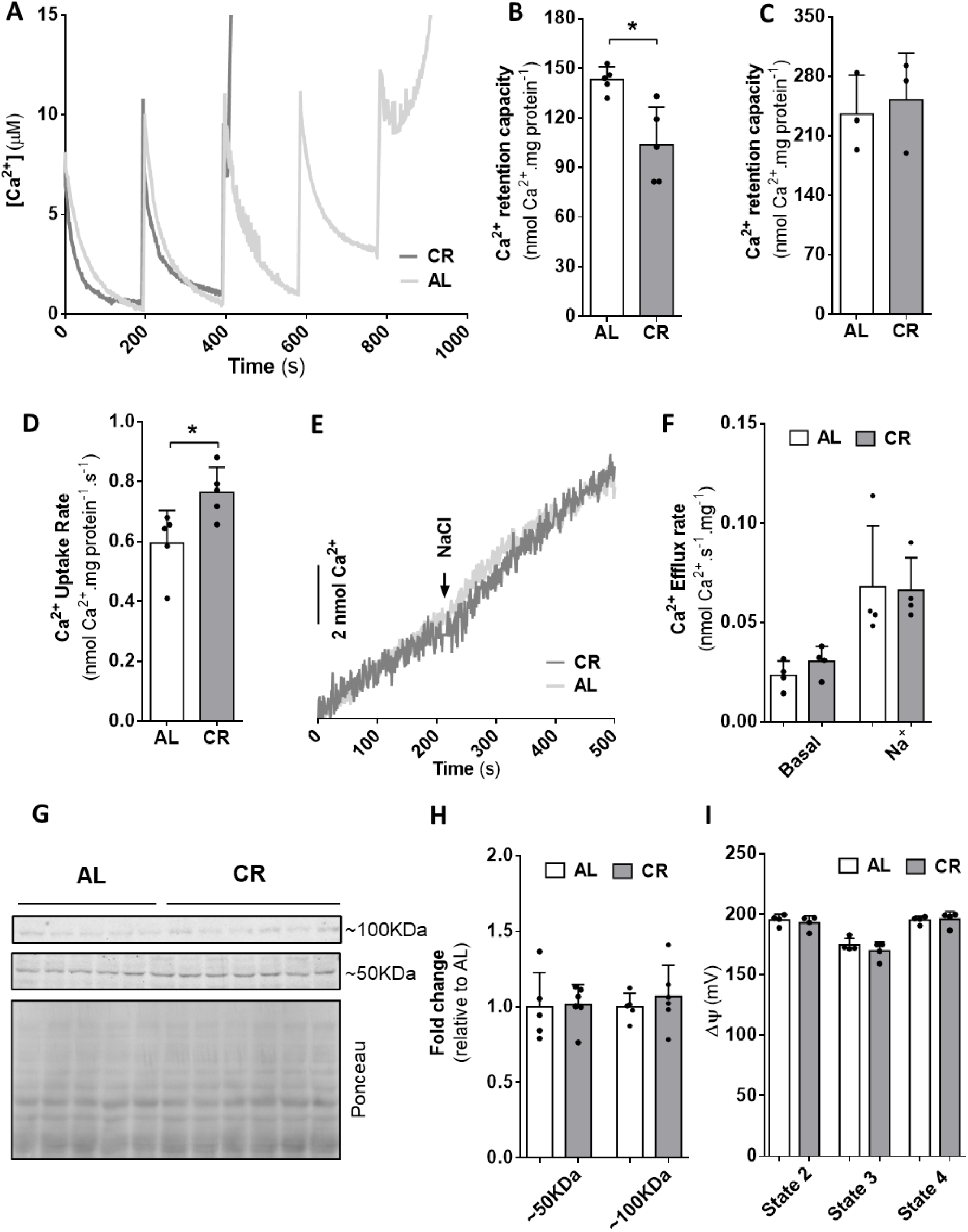
CR increases calcium uptake rates and sensitivity to mPT in kidney mitochondria. Mitochondria (500 μg protein) were incubated in 2 mL calcium uptake media, as described in the methodology. **A.** Representative calcium uptake traces for samples derived from AL (light lines) and CR (dark lines) kidneys; 10 μM calcium additions were made every 200 s. **B.** Maximum calcium retention capacity, quantified from traces such as in Panel A. **C.** Maximum calcium retention capacity in the presence of 10 μM cyclosporin A (CsA). **D.** Calcium uptake rates, quantified from traces such as in Panel A. **E.** Representative calcium efflux trace. The first part of the trace represents the basal mitochondrial calcium efflux, induced after addition of ruthenium red (RuR). Calcium efflux was stimulated with 20 mM NaCl, added where indicated. **F**. Calcium efflux rates were measured from traces such as in Panel E. **G.** Representative western blot images for NCLX and upload control Ponceau staining; ~100 KDa and ~50 KDa bands correspond to the NCLX dimer and monomer, respectively. **H.** Densitometric quantification for the blot signals, relative to AL. Protein was normalized in each lane by the total Ponceau stain. **I.** Mitochondrial membrane potentials were measured as described in Methods under state 2 (succinate + rotenone), state 3 (ADP), and state 4 (oligomycin) conditions. * p<0.05.

Interestingly, kidney mitochondria from CR animals also displayed significantly higher Ca^2+^ uptake rates (Fig. 3D), suggesting that the diet not only affects the response of these organelles to large, usually damaging, Ca^2+^ concentrations, but also alters physiological ion homeostasis in this organelle. Net calcium transport rates result from both influx and efflux rates. The increase in the net Ca^2+^ influx by CR could, therefore, be explained by decreased efflux. To measure Ca^2+^ efflux, mitochondria were loaded with Ca^2+^ and then the MCU was inhibited, resulting in Ca^2+^ efflux, which was measured both in the absence (Fig. 3E and F, “basal”) and presence of Na^+^, to stimulate Na^+^/Ca^2+^ exchange through NCLX (36). No changes in basal or Na^+^-induced mitochondrial calcium efflux were observed in CR versus AL samples. Indeed, NCLX quantities (Fig. 3G and H) were equal between CR and AL mitochondria; the protein was detected as both a monomer and dimer, at ~50 and ~100KDa (48).

Since enhanced Ca^2+^ uptake rates were not related to lower extrusion, and small but significant changes in respiration were detected (Fig. 2A), we quantitatively determined mitochondrial inner membrane potentials (ΔΨ), which are the driving force for Ca^2+^ uptake (49,50). No differences were observed between CR and AL mitochondria in any respiratory state (Fig 3I). Therefore, the changes in Ca^2+^ uptake rates measured in Fig 3D are not attributable to ΔΨ.

Changes in mitochondrial Ca^2+^ uptake may also be associated with altered quantities and structure of the MCU complex, which were assessed in Figure 4. No changes in the content of the MCU itself were observed. EMRE and MICU1 content were also unchanged in response to CR. However, MICU2 expression was approximately 25% lower. Since MICU2 expression is associated with lower Ca^2+^ uptake rates (16,17,51), its decrease may explain the higher calcium uptake rates observed in CR kidney mitochondria. Interestingly, this is the first evidence of diet-induced MICU2 modulation, to our knowledge. Given this potentially novel finding, we studied the MCU composition in mitochondria from two additional organs: heart and liver. Liver mitochondria from CR animals have higher Ca^2+^ uptake rates (30), and also exhibited lower MICU2 quantities in response to CR (Fig. 4C,D). MICU1 remained constant. On the other hand, heart mitochondria do not exhibit changes in Ca^2+^ uptake rates with CR (31), and also had no change in MCU, MICU1 or MICU2 complex (Fig. 4E,F). Taken together, this demonstrates that MICU2 is regulated by CR in an organ-specific manner, which correlates with the higher calcium uptake rates observed in kidney and liver.

**Figure 4.**
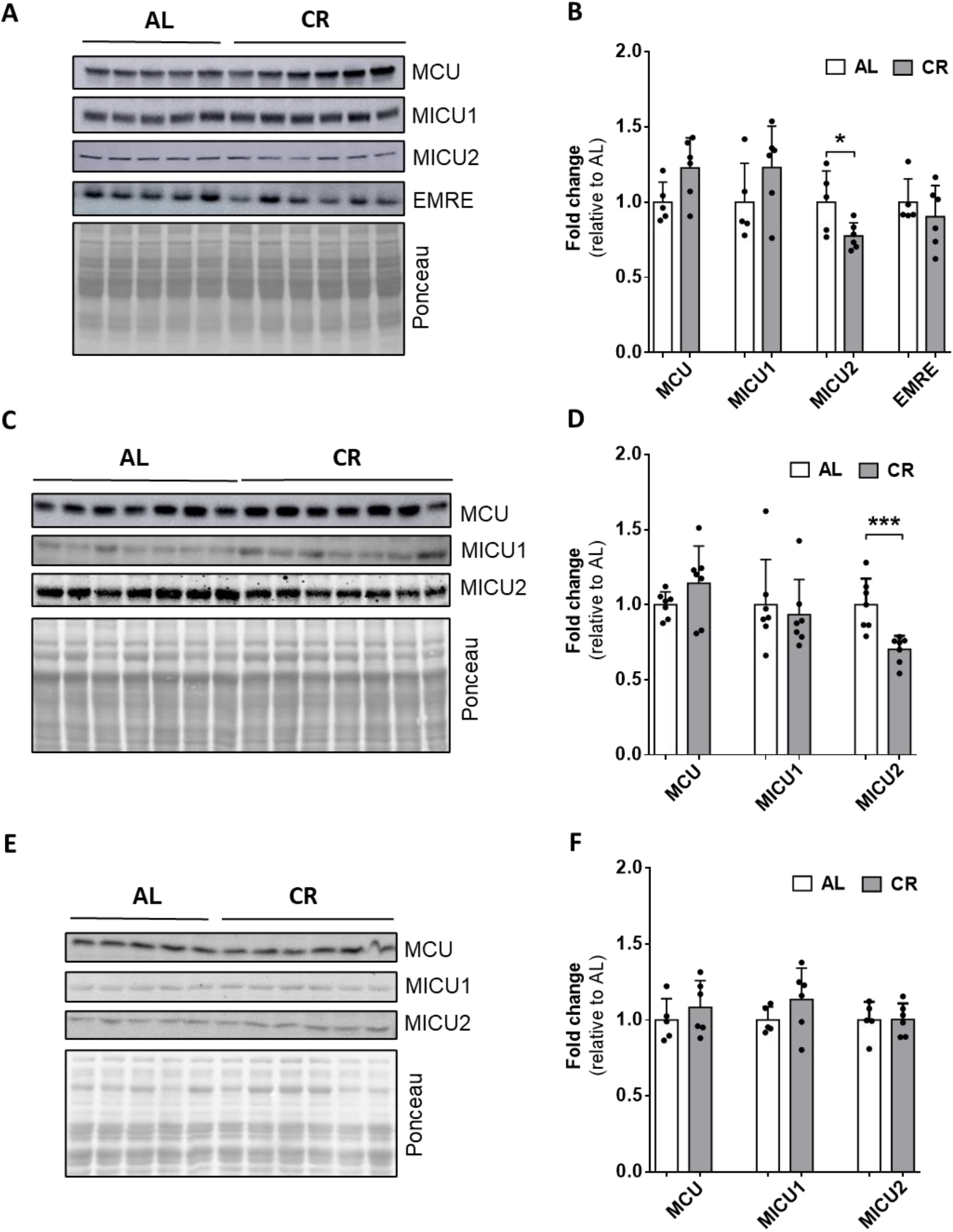
CR decreases MICU2 levels in kidney and liver isolated mitochondria. **A**. Representative western blot of the MCU complex in kidney mitochondria: MCU, MICU1, MICU2 and EMRE and loading control Ponceau stained images. Representative western blot and Ponceau staining images for liver **(C)** and heart **(E)** mitochondria. Densitometric analysis for the blot images (relative to AL) for kidney **(B)**, liver **(D)** and heart **(F)** mitochondria; protein signal was normalized by the total Ponceau staining signal of each lane. Light gray and dark gray bars represent samples from AL and CR rats. *p < 0.05 and ***p < 0.001.

While the decrease in MICU2 mediated by CR in kidney explains higher Ca^2+^ uptake rates, it is not directly linked to modulation of the mPT. Prior data in liver mitochondria indicate that Ca^2+^ retention capacity and uptake rates may have an inverse relationship (52). We thus investigated (Figure 5) if the higher Ca^2+^ uptake rates in kidney mitochondria were the cause for mPT and limited uptake capacity in kidney CR samples. To do this, we measured uptake rate and capacity in control mitochondria using three different concentrations of Ca^2+^ additions (Fig. 5A). We found that lower Ca^2+^ boluses were associated with lower uptake rates (Fig. 5B). Interestingly, lower Ca^2+^ uptake rates in kidney mitochondria decreased maximum uptake capacity (Fig. 5C). This demonstrates that the enhanced sensitivity to mPT and low maximum Ca^2+^ uptake capacity promoted by CR are not a consequence of MICU2 deficiency-induced accelerated Ca^2+^ transport.

**Figure 5.**
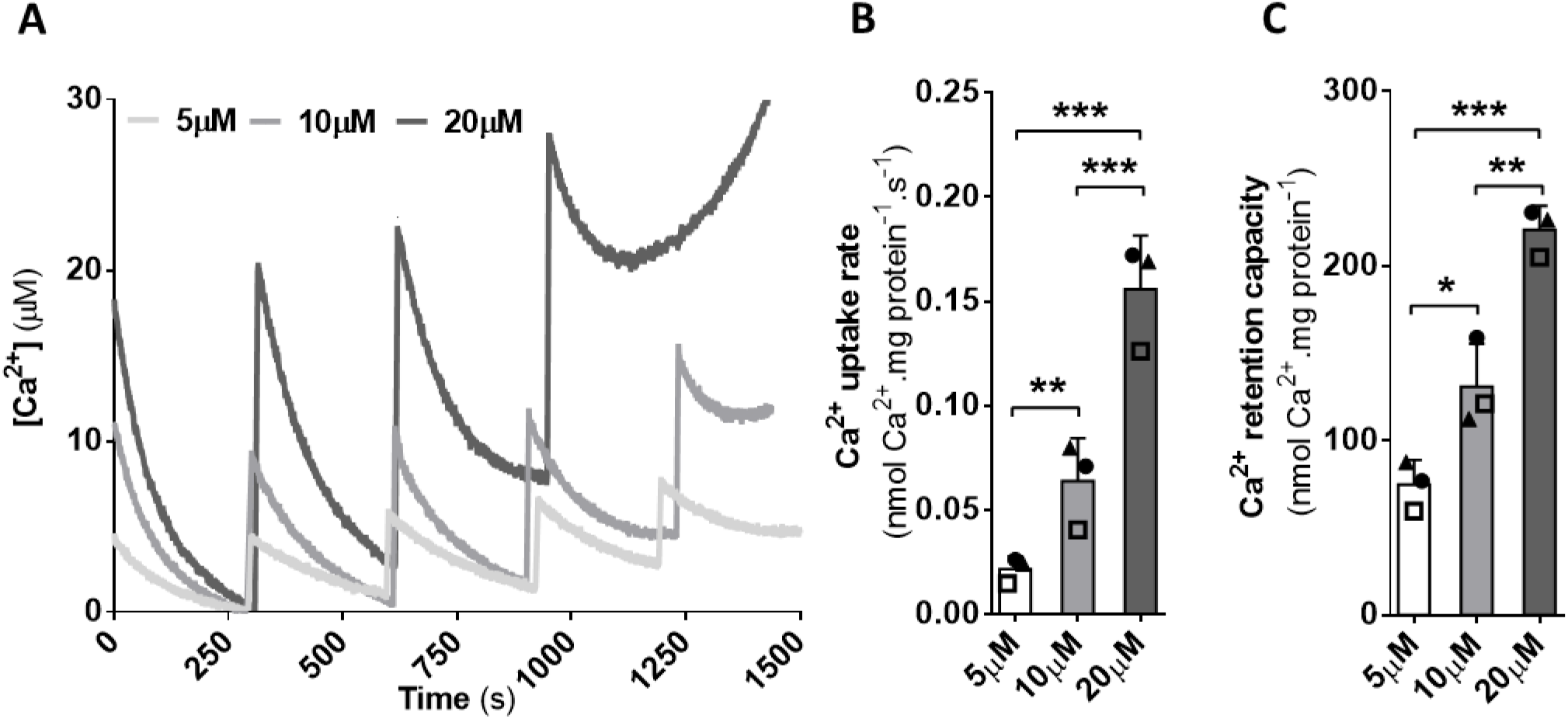
Ca^2+^ retention capacity increases with higher uptake rates. Calcium uptake was measured as described in the methodology, using sequential additions of 5 μM, 10 μM and 20 μM Ca^2+^, as indicated in Panel A. **B.** Calcium uptake rates. **C.** Calcium retention capacity. *p < 0.05, **p < 0.01 and ***p < 0.001.

Other mitochondrial properties previously related to CR and mPT modulation are changes in acetylation levels mediated by deacetylase Sirt3 (29) and changes in mitochondrial morphology (53). We thus probed for changes in Sirt3 expression and modulation of the quantities of proteins involved in mitochondrial morphology (Mnf2, Opa1, Oma1, and Drp1; Supplementary Figure 1), but found no effect of CR on these markers. Thus, changes in acetylation and mitochondrial morphology cannot be related to the enhanced mPT observed in CR kidney mitochondria.

Changes in redox state are also widely associated with CR and involved in mPT; while CR prevents oxidative damage in many tissues (54), a more oxidized redox state is associated with opening of the mPT pore (8). We thus evaluated redox state in CR kidneys. Overall redox state in kidney mitochondria was not significantly changed, as assessed by levels of carbonylated proteins and methionine sulfoxide (Supplementary Figure 2S). We also determined that important mitochondrial antioxidants including nicotinamide nucleotide transhydrogenase (NNT, which provides most mitochondrial NADPH), mitochondrial manganese-dependent superoxide dismutase (SOD2) and uncoupling protein (UCP2, which controls membrane potentials and, hence, electron transport and leakage) were not differently expressed in CR versus AL mitochondria (Fig. 6 A and B). On the other hand, direct measurement of H_2_O_2_ released from mitochondria showed a ~50% increase in CR mitochondria compared to AL. This increase may be related to the enhanced oxygen consumption rates (Fig. 2A) in the presence of equal ΔΨ (Fig. 3I) and could cause enhanced mPT (Fig. 3A,B), which is widely associated with oxidative imbalance (8,23).

**Figure 6.**
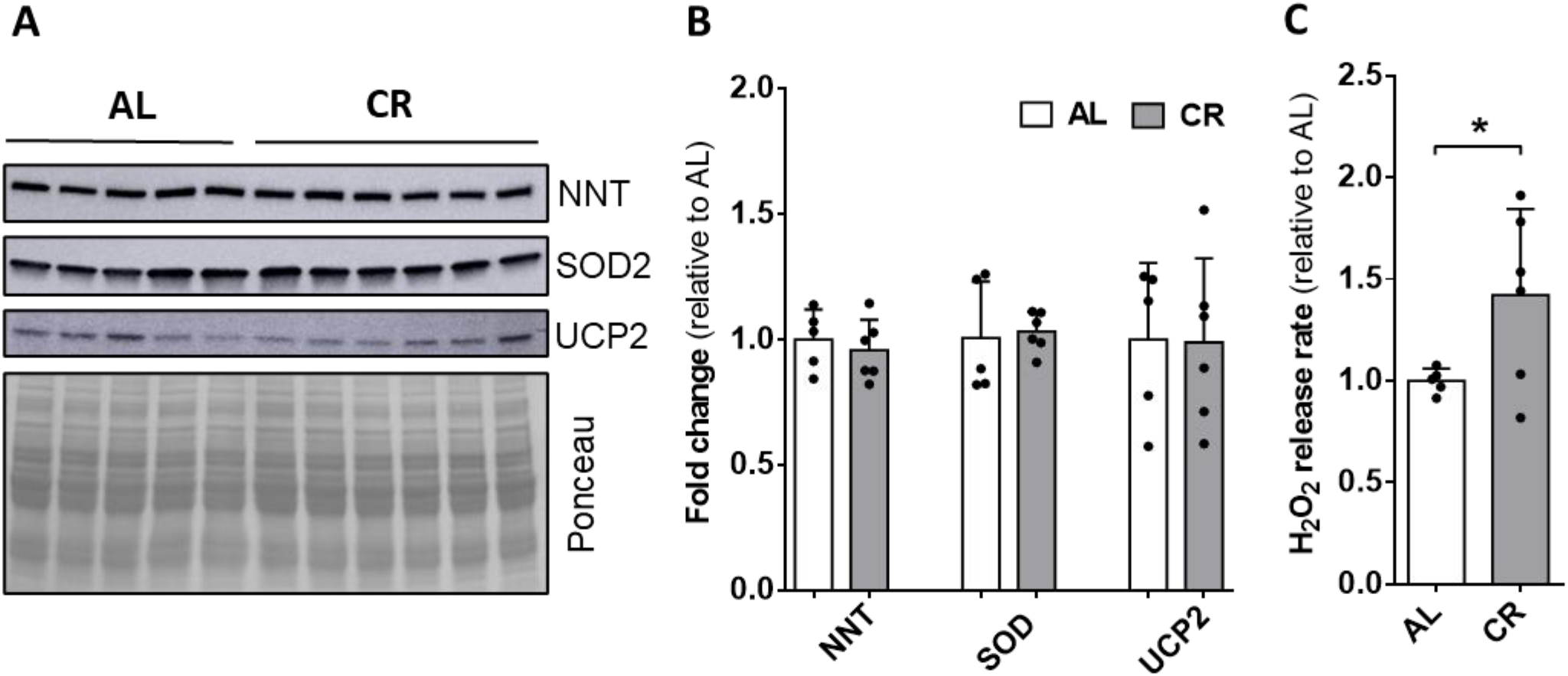
CR increases mitochondrial peroxide efflux rates without changes in mitochondrial antioxidant systems. Protein levels were assessed by western blot analysis in isolated mitochondrial samples. **A.** Representative western blot images for proteins involved in oxidant clearance: NNT, SOD2 and UCP2. **B.** Densitometry analysis for each blot image (relative to AL); protein signal was normalized by the total Ponceau staining signal of each lane. **C.** Peroxide release rates were measured under state 2 using the Amplex Red/Horseradish peroxidase system, as described in the methodology. Light gray and gray bars represent samples of mitochondria from AL and CR rats *p < 0.05.

## Discussion

CR is an intervention well known to enhance lifespans and healthspans, preventing many age-related diseases. The biochemical mechanisms in which increased health is promoted by CR are still being uncovered, and we, as well as other authors, have found that its metabolic impacts are tissue-specific (54,55). A novel metabolic change promoted by CR is altered mitochondrial Ca^2+^ uptake (28–31). As calcium ions are central metabolic regulators both in mitochondria and the cytosol, their organellar uptake properties are certainly impactful for the outcome of the interventions that change them. Indeed, prior work from our lab has shown that changes in mitochondrial Ca^2+^ uptake in liver and brain lead to protection against ischemic and excitotoxic damage (28–30).

Here, we demonstrate that CR increases mitochondrial calcium uptake rates also in the kidney, without modifying efflux activity (Fig. 3). These CR-promoted alterations in the kinetics of calcium transport are of interest from a physiological point of view, as matrix calcium regulates several aspects of mitochondrial function, including oxidative phosphorylation and redox balance (5,6,8,56,57). Higher uptake rates lead to stronger calcium transients in the matrix, and will make these organelles more responsive to cytoplasmic calcium signals of the same magnitude (16,17,21,51). The kinetics of Ca^2+^ transport in CR kidney mitochondria are also consistent with higher basal matrix concentrations. We hypothesize that these basal matrix Ca^2+^ concentrations in the CR group may be responsible for the higher oxygen consumption rates under the respiratory states 2, 4 and 3U (Fig. 2), as well as higher H_2_O_2_ efflux rates (Fig. 6). This is supported by the lack of alterations in the amount or activity of electron transport complexes.

Higher mitochondrial Ca^2+^ uptake rates in the CR group were not due to altered inner membrane potentials as a driving force, nor in changes in the expression of NCLX or the MCU (50; Figs. 3 and 4). Since other proteins in the MCU complex are important to determine activity (9), we checked for quantities of these components, and found that MICU2 was downregulated by CR (Fig. 4). Interestingly, in liver mitochondria, but not heart, MICU2 was also downregulated, suggesting that MICU2 is a diet-regulated MCU complex component that modulates mitochondrial Ca^2+^ uptake in a tissue-dependent manner.

Indeed, higher MICU1/MICU2 ratios promote enhanced mitochondrial Ca^2+^ uptake rates and higher basal matrix Ca^2+^ concentrations (16,17,51). Furthermore, in MICU2-silenced cells, mitochondrial Ca^2+^ uptake is increased, leading to lower intensity cytoplasmic Ca^2+^ signals (16). Under these conditions, MICU1 is upregulated and forms homodimers. Avoiding MICU2 interaction with MICU1 (MICU1 C463A mutant) increases mitochondrial Ca^2+^ uptake, as reflected by longer and higher intensity mitochondrial Ca^2+^ transients (17). In our system, the lower amount of MICU2 was probably not associated to homodimerization of MICU1, as no changes in total MICU1 were detected. An MCU complex without MICU2 is completely functional and exhibits higher Ca^2+^ uptake ability. Since CR promotes an increase in the content of MICU2-deficient MCU complexes in rat kidney and liver, the molecular signature is consistent with the higher Ca^2+^ uptake rates we measured.

This is the first description, to our knowledge, that MICU2 can be regulated by diet. Several works support the idea that the activity of the MCU complex is regulated through alterations in MICU1/2 protein expression under pathological contexts. In diabetic mouse hearts, MICU1 is downregulated, contributing toward cardiomyocyte cell death (19). Polycystin 2 knockdown also induces MICU2 downregulation, favoring mitochondrial calcium uptake (21). Detrimental effects are found in hearts (58) and the nervous system (59) as a result of a deficient function/amount of MICU2. In several cancer models, mitochondrial Ca^2+^ transport is impaired through alterations in the composition of MICU proteins (20). Our results add, for the first time, a physiological context in which MICU2 is downregulated, in response to a systemic decrease in caloric intake.

Interestingly, while kidney mitochondria respond to CR with increased Ca^2+^ uptake rates, as seen in liver and brain (29,30), they differ significantly from these tissues when maximum uptake capacity (limited by mPT) is compared (Fig. 3). This could indicate that, contrary to the mostly protective effects of CR, it may make kidneys more prone to damaging conditions which increase Ca^2+^ intracellularly and can promote mPT. No studies have looked at kidney resilience in long-term CR to answer this point. However, short-term total food restriction and fasting protect kidneys against ischemia/reperfusion damage (32). Since the effect of these nutritional interventions is mimicked by CsA, they suggest decreased mitochondrial sensitivity to Ca^2+^-mPT (32), although no direct proof of this mechanism was provided. The reasons for these differences may be many, including other mechanisms protecting against ischemia/reperfusion in the kidneys, as well as the different duration and type of dietary interventions (33). Further studies should explore this point, as our results suggest it is plausible that long-term CR leads to kidney vulnerability to damage induced by excessive intracellular Ca^2+^.

We sought to explore the mechanisms of enhanced mPT in CR kidneys here, and found that CR also induces small changes in respiratory rates and increases H_2_O_2_ release from mitochondria. Once again, this is an unexpected result, since in most tissues CR decreases H_2_O_2_ release and improves redox state (54). Redox alterations in the mitochondrial environment are determinant for mPT susceptibility; oxidants promote protein thiol oxidation and favor mPT induction (8,23). Previous work demonstrates that CR increases the activity/amount of several antioxidant systems in rodent kidneys, including the activity/content of catalase (54). This is compatible with the higher levels of H_2_O_2_ release we see from mitochondria, but preserved overall cellular redox state (Figs. S2 and 6). It is tempting to hypothesize that retrograde mitochondrial redox signals promoted by H_2_O_2_ may modulate cell responses to cope with oxidant stresses, including the expression of cellular antioxidants.

Altogether, we show that CR modulates calcium uptake in kidney mitochondria. This involves higher uptake rates and lower resilience against Ca^2+^-mPT. At the molecular level, these changes were consistent with an increase in the content of MICU2-deficient MCU complexes, as well as enhanced H_2_O_2_ generation by CR mitochondria. Both the loss in Ca^2+^ uptake capacity and enhanced oxidant production by CR kidney mitochondria are unexpected, given their association with damaging conditions, while CR is mostly considered a protective dietary intervention. This uncovers the need for more focused and tissue-specific studies uncovering the metabolic results of restricted diets.

## Acknowledgements

This work was supported by *Conselho Nacional de Desenvolvimento Científico e Tecnológico* (CNPq), *Coordenação de Aperfeiçoamento de Pessoal de Nível Superior* (CAPES) finance code 001 and *Fundação de Amparo à Pesquisa do Estado de São Paulo* (FAPESP) grants 20/06970-5, 19/04233-6, and 19/05226-3, as well as FAPESP *Centro de Pesquisa, Inovação e Difusão de Processos Redox em Biomedicina (Redoxoma*) 13/07937-8. The authors are in debt to Silvânia M. P. Neves and her animal facility crew for expert animal care.

## Figure Legends

**Figure 1S.**
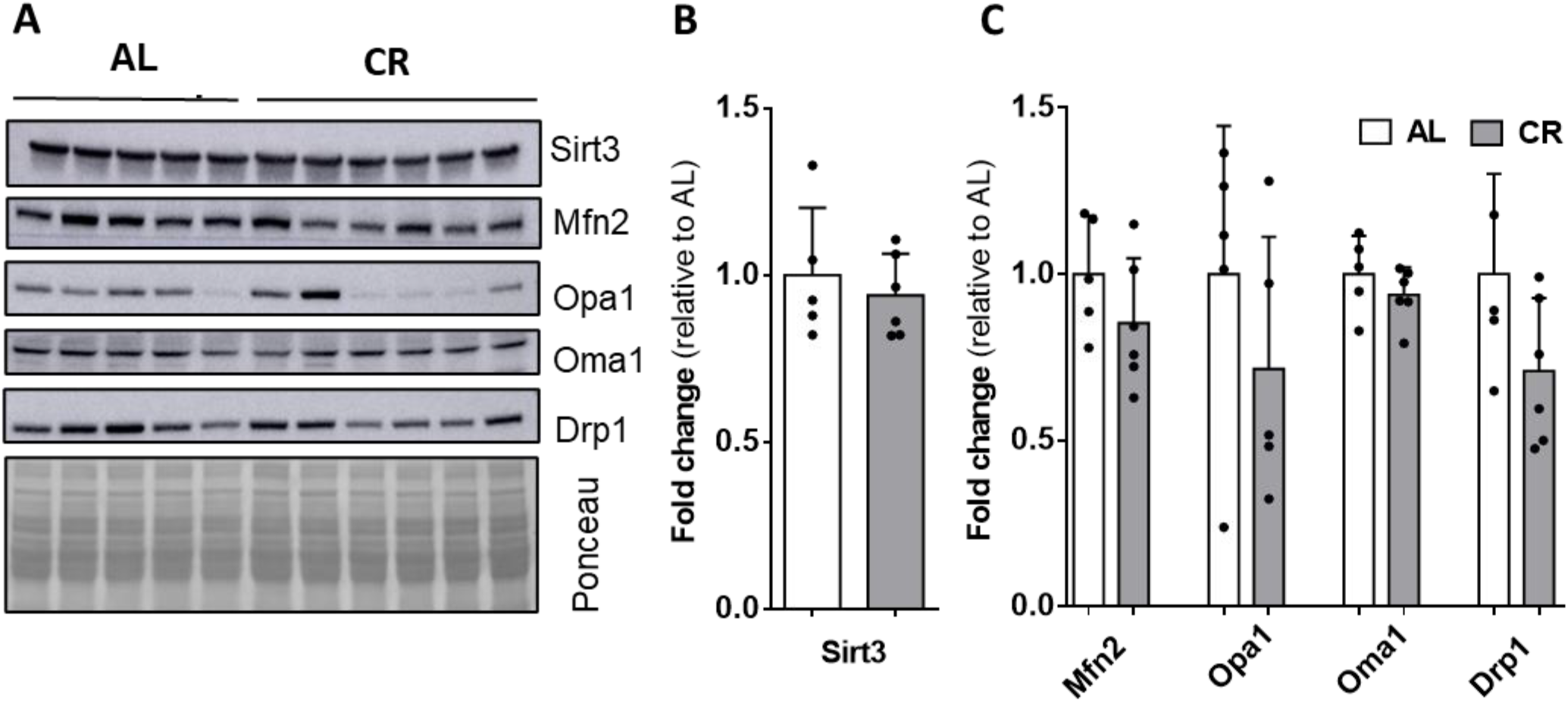
CR does not promote changes in the levels of proteins involved in the control of kidney mitochondrial morphology or acetylation. Protein levels were measured by western blot in isolated mitochondrial samples. **A.** Representative blot images for proteins involved in the control of mitochondrial acetylation (Sirt3) or morphology (Drp1, Mfn2, Opa1, Oma1). Densitometric analysis (relative to AL) of Sirt3 **(B)** and morphology proteins **(C)**. Protein signal was normalized by the total Ponceau staining signal of each lane. Light and dark bars represent samples of mitochondria from AL and CR kidney mitochondria.

**Figure 2S.**
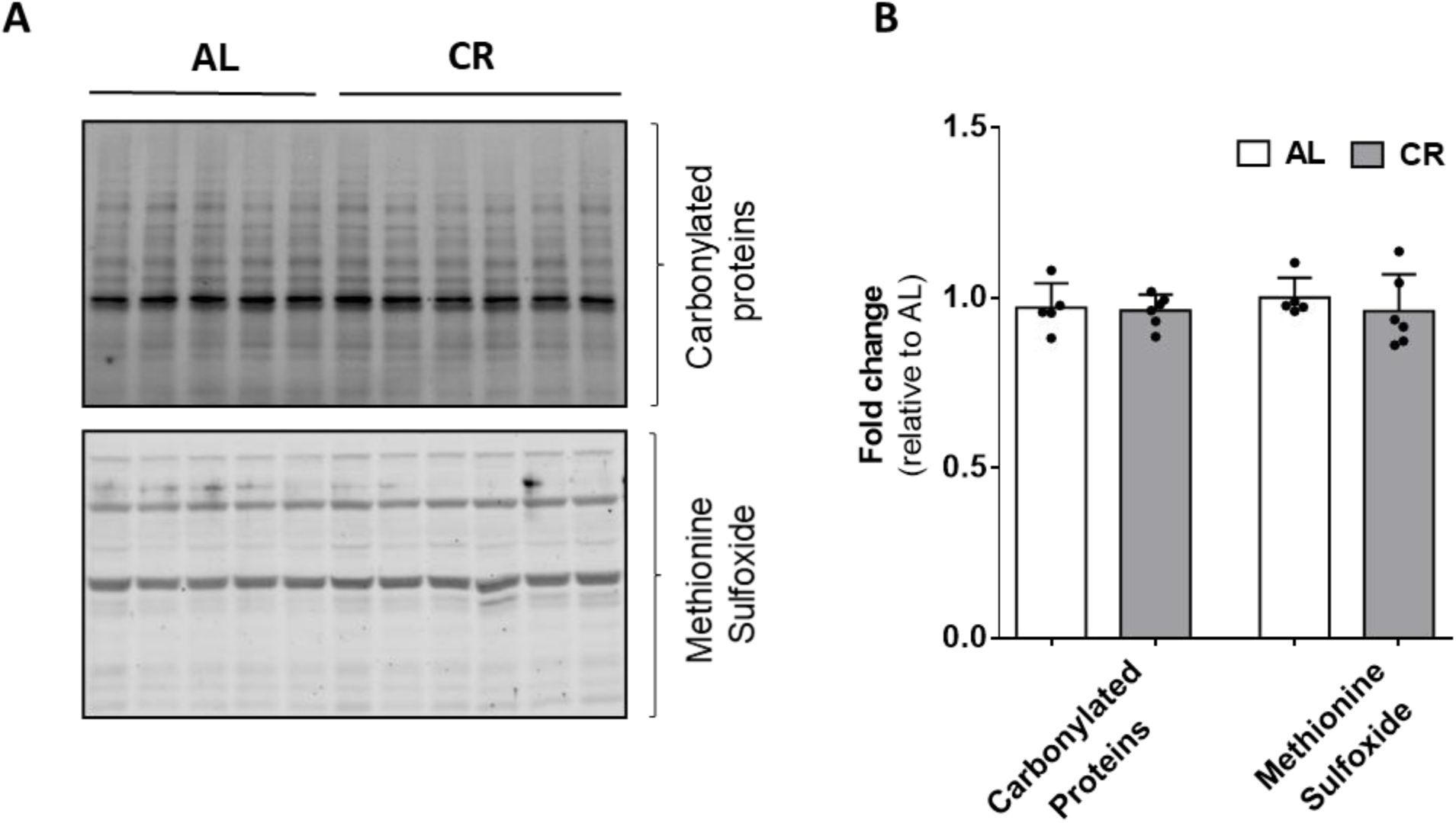
CR does not alter carbonylated protein levels or methionine sulfoxide in isolated mitochondria. Protein levels were assessed by western blot analysis in isolated mitochondrial samples. **A.** Representative blot images for total carbonylated proteins and methione sulfoxide. **B.** Densitometric analysis. Protein signal was normalized by the total Ponceau stain. Light and dark bars represent samples of mitochondria from AL and CR kidney mitochondria.

